# Parameter Efficient Fine-tuning of Transformer-based Masked Autoencoder Enhances Resource Constrained Neuroimage Analysis

**DOI:** 10.1101/2025.02.15.638442

**Authors:** Nikhil J. Dhinagar, Saket S. Ozarkar, Ketaki U. Buwa, Sophia I. Thomopoulos, Conor Owens-Walton, Emily Laltoo, Chirag Jagad, Yao-Liang Chen, Philip Cook, Corey McMillan, Chih-Chien Tsai, J-J Wang, Yih-Ru Wu, Paul M. Thompson

## Abstract

Recent innovations in artificial intelligence (AI) have increasingly focused on large-scale foundational models that are more general purpose in contrast to conventional models trained to perform specialized tasks. Transformer-based architectures have become the standard backbone in foundation models across data modalities (image, text, audio, video). There has been a keen interest in applying parameter-efficient fine-tuning (PEFT) methods to adapt these models to specialized downstream tasks in language and vision. These methods are particularly essential for medical image analysis where the limited availability of training data could lead to overfitting. In this work, we evaluated different types of PEFT methods on pre-trained vision transformers relative to typical training approaches, such as full fine-tuning and training from scratch. We used a transformer-based masked autoencoder (MAE) framework, to pretrain a vision encoder on T1- weighted (T1-w) brain MRIs. The pretrained vision transformers were then fine-tuned using different PEFT methods that reduced the trainable model parameters to as few as 0.04% of the original model size. Our study shows that: 1. PEFT methods were competitive with or outperformed the reference full fine-tuning approach and outperformed training from scratch, with only a fraction of the trainable parameters; 2. PEFT methods with a 32% reduction in model size boosted Alzheimer’s disease (AD) classification by 3% relative to full fine-tuning and 11% relative to a 3D CNN, with only 258 training scans; and 3. PEFT methods performed well on diverse neuroimaging tasks including AD and Parkinson’s disease (PD) classification, and “brain-age” prediction based on T1-w MRI datasets - a standard benchmark for deep learning models in neuroimaging; 4. smaller model sizes were competitive with larger models in test performance. Our results show the value of adapting foundation models to neuroimaging tasks efficiently and effectively in contrast to training stand- alone special purpose models.

## 1. INTRODUCTION

Recent advances in AI have greatly advanced medical image analysis. AI techniques can be used in clinical practice as well as for neuroscience and radiology research. Many promising studies have trained deep learning methods on brain magnetic resonance imaging (MRI) data collected worldwide to perform tasks such as differential diagnosis or prognosis in dementia. Some key challenges in adapting AI to medical tasks include working with limited labeled training data, privacy requirements when handling medical data, and heterogeneity of data collected at different sites and scanners. The ‘transfer learning’ paradigm, which pre-trains AI models first and then fine-tunes them for specific tasks, has helped to overcome the lack of large datasets in medical applications. Many pre-training approaches have been proposed and tested, including methods based on (semi-) supervised learning and contrastive learning.^1 2 3 4^ Masked autoencoders provide an effective way to pretrain vision transformers. ^5 6^

The pre-trained models are then adapted to a downstream task by fully fine-tuning all the neural network layers or some of the layers. As large-scale foundation models have emerged, ^7^ methods are needed to efficiently fine-tune them. The use of large vision transformer-based models is challenging, as they are prone to overfitting with smaller datasets. Parameter-efficient fine-tuning (PEFT) methods have seen increased interest in the context of language models before being adopted for computer vision applications as well. PEFT methods only fine-tune a small fraction of the model parameters and freeze the remainder. The tunable model parameters in the PEFT methods are usually modified through *additive* or *selective* approaches applied to different sections and mechanisms of the model. PEFT methods also make it possible to fine-tune larger models on limited computational consumer hardware. PEFT methods include adaptors, ^8 9^ low-rank approximation (LORA), ^10^ prompt tuning, ^11^ and attention tuning.^12^ Some recent work shows the benefits of PEFT methods for medical applications. ^13 14^ For some tasks, PEFT may perform as well as full fine-tuning, at a fraction of the computational cost. In this work, we evaluated the performance of different *additive* and *selective* PEFT methods with the vision transformer. We tested the effect of PEFT methods on the sample size requirements for the downstream fine-tuning dataset. We also demonstrated that these methods can be used to adapt the vision encoders to diverse neuroimaging tasks and their corresponding datasets.

## 2. METHODS

In this work, we used the vision transformer (ViT) architecture, ^15^ specifically, the ViT-Base (ViT-B), ^16^ and its smaller variant, ViT-Small (ViT-S). Each of these models is based on the transformer architecture, currently used across diverse data modalities (image, video, audio, text) ubiquitously. The ViT-B encoder has 88 million trainable parameters, and the ViT-S encoder has 12 million trainable parameters. We pretrained the ViT encoders on the large-scale UK Biobank (UKBB) brain MRI datasets using the MAE framework. ^5 6^ 75% of patches generated from the T1-w MRIs were masked initially and a lightweight decoder was used to predict the masked patches.

Given the state-of-the art nature of this framework and relatively large, diverse pre-training data, this foundation model is expected to *generalize* well to downstream tasks. In the next section, we describe PEFT methods that enabled this foundation model to be further *adapted* to achieve the best performance possible on specialized tasks and their datasets. In this paper, we evaluated two main types of PEFT methods, shown in **Table 1**: *additive* methods that require inclusion of new parameters and *selective* methods that optimize only a subset of the model’s parameters. In both methods, the underlying original model parameters are frozen and only the new or subset parameters are fine-tuned. As an *additive* method we tested low-rank adaptation (LORA), ^10 17^ where the pre-trained attention weight matrix is frozen and trainable rank decomposition matrices are injected in the transformer encoder layers. The *selective* methods consisted of two approaches, attention fine-tune, and layer normalization fine-tune, where only the attention and layer norm layers are respectively fine-tuned. As a baseline, we compared our PEFT methods with the encoders trained from scratch (without any pre-training) and full fine-tuning involving all the original trainable model parameters. We also used a 3D DenseNet121 CNN with 11 million parameters as an additional reference for our experiments.

**Table 1.** Model Parameter count (in millions) using parameter-efficient fine-tuning methods with the ViT-B and ViT-S.

| Backbone | Scratch | Full fine-tune | Attention fine-tune | LORA | Layernorm fine-tune |
| --- | --- | --- | --- | --- | --- |
| ViT-B | 88,204,034 | 88,204,034<br>(100%) | 28,349,954<br>(32%) | 1,181,308<br>(1.3%) | 38,400<br>(0.04%) |
| ViT-S | 12,220,800 | 12,220,800<br>(100%) | 3,548,930<br>(29%) | 295,804<br>(2.4%) | 10,754<br>(0.09%) |

## 3. EXPERIMENTAL SETUP

### 3.1 Data

In line with similar studies, ^2^ all 3D T1-weighted brain MRI scans were pre-processed via standard steps for neuroimaging analyses, including: nonparametric intensity normalization (N4 bias field correction), ^18^ ‘skull-stripping’, linear registration to a template with 9 degrees of freedom, and isotropic resampling of voxels to 2-mm resolution. The input spatial dimension of the MRIs was 91×109×91. All images were *z*-transformed (setting each image’s mean and SD to a standard value) to stabilize model training. The T1-w scans were re-sized to 80 voxels across all dimensions before model training. In this work, we also used 3D T1-w brain MRI scans from the UK Biobank ^19^ dataset to pretrain the vision encoders using the MAE framework (see Table 2 for details). We used data from the publicly available ADNI dataset ^20^ for the downstream tasks of Alzheimer’s disease (AD) classification and brain age prediction. We used two additional datasets for Parkinson’s disease (PD) classification. We also used the OASIS ^21^ and UPenn datasets as out-of-distribution datasets for zero-shot testing for each experiment. For the brain age experiments, we used a subset of ADNI with only MRIs from healthy control participants. The datasets used in this study are summarized in **Table 2**.

**Table 2.** Summary of data utilized in this work for pretraining and fine-tuning.

| Dataset | N | #Subjects | Age (years) | Sex (F/M) |
| --- | --- | --- | --- | --- |
| UKB | 38,703 | 38,703 | 44.6-82.8 | 20,216F/18,487M |
| ADNI | 4,098 | 1,188 | 55.7-92.8 | 556F/632M |
| OASIS | 600 | 600 | 43.5-97.0 | 341F/359M |
| Taiwan | 467 | 467 | 20.0-80.0 | 236F/231M |
| UPenn | 164 | 164 | 50.0-86.0 | 63F/101M |

### 3.2 Model Training and Testing

#### Dataset

We performed a random search to select hyperparameter values including the batch size, learning rate, learning rate scheduler warmup epochs, weight decay, and dropout for pretraining and finetuning. The vision transformers in the MAE had the following specifications, ViT-B - Encoder: 12 layers, 12 attention heads and Decoder: 8 layers, 12 attention heads; and ViT-S – Encoder: 6 layers, 8 attention heads and Decoder: 8 attention heads, 4 layers. A patch size of 16 was used for the ViTs. The ViTs were pre-trained for 1,000 epochs and fine-tuned for 30 to 50 epochs with an early stopping of 10 epochs. All models used the AdamW optimizer and minimized the mean squared error (MSE) loss for pre-training, the L1, binary cross entropy loss functions for regression and classification fine-tuning tasks respectively. We evaluated our models with the receiver-operator characteristic curve-area under the curve (ROC-AUC), and mean absolute error (MAE) for classification and regression, respectively. In our results, we present an average over multiple runs with three different seeds. We conducted zero-shot testing with an independent dataset for each of the fine-tuning tasks.

## 4. RESULTS

The ViTs were trained from scratch and fully fine-tuned as a baseline. We evaluated the PEFT methods relative to training from scratch, with full fine-tuning -- as well as the 3D DenseNet121 CNN. We investigated the use of PEFT methods for different neuroimaging tasks and datasets as presented in **Tables 3 and 4. Figure 1** Illustrates the tradeoff between test performance and the number of model parameters; **Figures 2 and 3** shows the test ROC-AUC versus training data sizes using ADNI, where the different curves represent the various PEFT methods.

**Table 3.** Application of PEFT methods using the ViT-B for diverse tasks, relative to full fine-tuning.

| Backbone |  | ViT-B |  |  |  |  | DenseNet121 CNN |
| --- | --- | --- | --- | --- | --- | --- | --- |
| Task | PEFT | Scratch | Full fine-tune | Attention fine-tune | LORA | Layernorm fine-tune | Scratch |
|  | Dataset |  |  |  |  |  |  |
| Alzheimer's Disease (AD) Classification, Test ROC-AUC | ADNI | 0.8531<br>(0.0304) | **0.8619**<br>**(0.0241)** | **0.8613**<br>**(0.0094)** | 0.8153<br>(0.0500) | 0.8077<br>(0.0269) | 0.8865<br>(0.013) |
|  | OASIS (zero-shot) | 0.8184<br>(0.0040) | 0.8353<br>(0.0103) | 0.8311<br>(0.0075) | 0.8295<br>(0.0162) | 0.8263<br>(0.0107) | 0.8371<br>(0.0080) |
| Brain Age Prediction, Test MAE | ADNI | 4.9140<br>(0.3700) | **4.5076**<br>**(0.1530)** | **4.0860**<br>**(0.1400)** | **4.4836**<br>**(0.1900)** | **4.2463**<br>**(0.5070)** | 4.4255<br>(0.0212) |
|  | OASIS (zero-shot) | 9.6226<br>(1.0190) | 8.0880<br>(0.9810) | 6.7593<br>(0.4060) | 8.4426<br>(0.3710) | 7.6873<br>(1.4000) | 7.8484<br>(0.2870) |
| Parkinson's Disease (PD) classification, Test ROC-AUC | Taiwan | 0.5424<br>(0.0927) | **0.6521**<br>**(0.0400)** | **0.6218**<br>**(0.1073)** | **0.6200**<br>**(0.0109)** | 0.5962<br>(0.0281) | 0.5891<br>(0.0263) |
|  | UPenn (zero-shot) | 0.4791<br>(0.0873) | 0.5457<br>(0.0325) | 0.5414<br>(0.1170) | 0.5710<br>(0.0019) | 0.5687<br>(0.0648) | 0.6100<br>(0.0393) |

**Table 4.** Application of PEFT methods using the ViT-S for AD classification, relative to full fine-tuning.

| Backbone |  | ViT-S |  |  |  |  |
| --- | --- | --- | --- | --- | --- | --- |
| Task | PEFT | Scratch | Full fine-tune | Attention fine-tune | LORA | Layernorm fine-tune |
|  | Dataset |  |  |  |  |  |
| Alzheimer's Disease (AD) Classification, Test ROC-AUC | ADNI | 0.8426<br>(0.0189) | **0.8766**<br>**(0.0092)** | **0.8633**<br>**(0.0101)** | 0.7886<br>(0.0267) | 0.7987<br>(0.0648) |
|  | OASIS (zero-shot) | 0.8316<br>(0.0053) | 0.8442<br>(0.0087) | 0.8306<br>(0.0163) | 0.8198<br>(0.0078) | 0.8122<br>(0.0417) |

**Figure 1.**
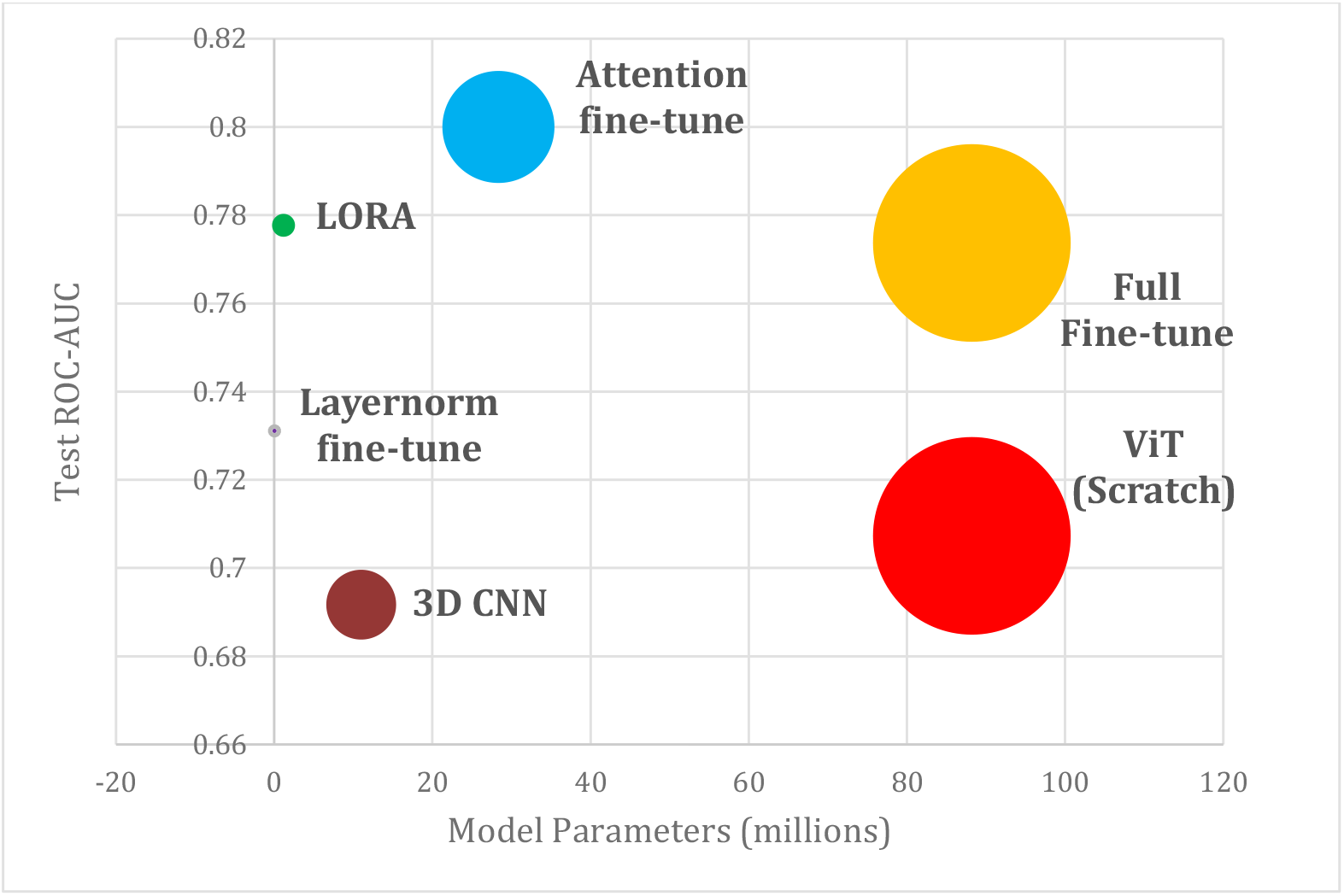
Test performance and vision transformer (ViT-B) model parameter tradeoff using different PEFT methods relative to full fine-tuning for AD classification - with *only 258 training MRIs* from the ADNI dataset. Larger bubble sizes indicate a greater number of model parameters to be tuned for the downstream task. Compared to very large models (on the right), attention fine-tuned and LORA models maintain high accuracy with far fewer parameters (top left).

**Figure 2.**
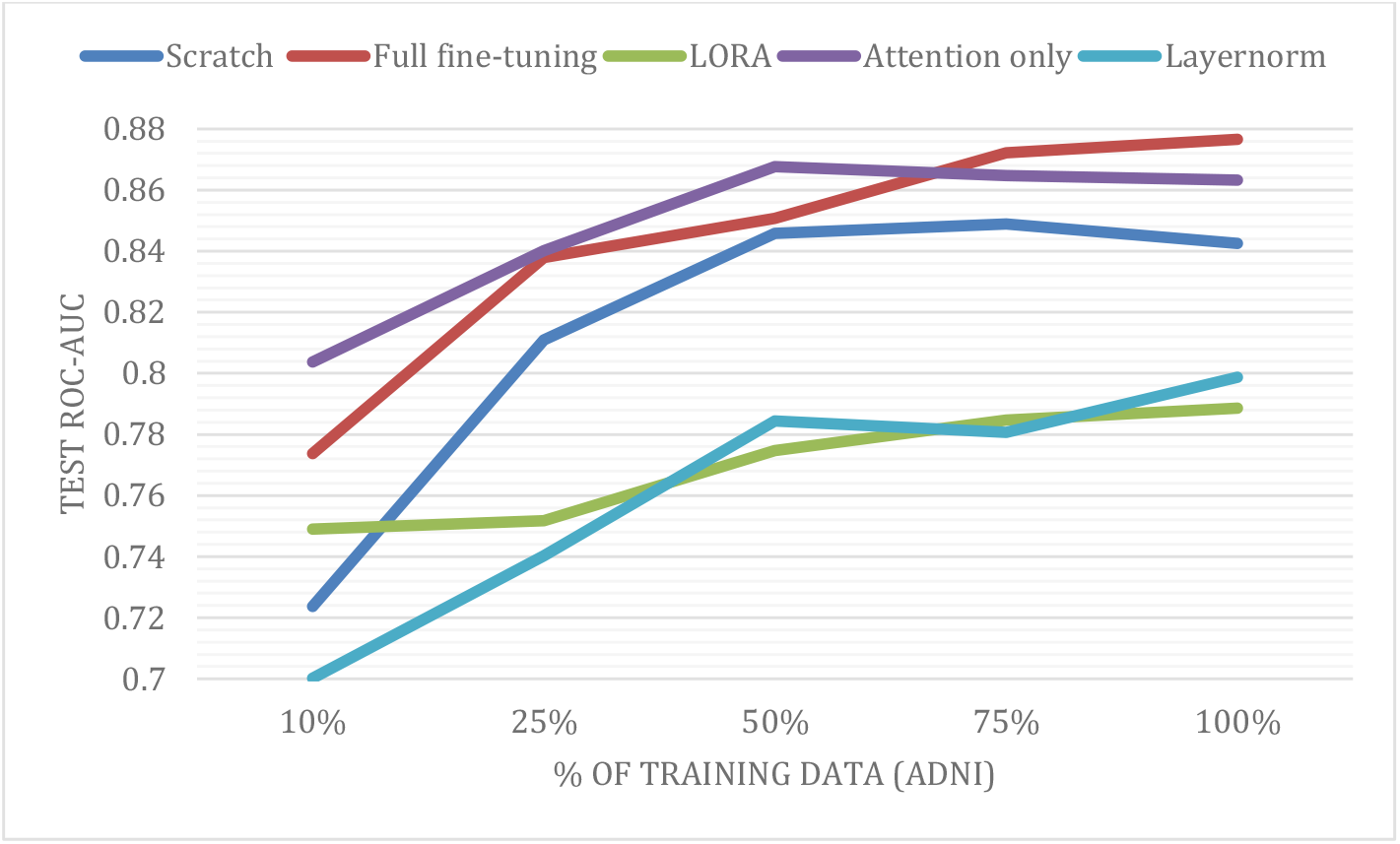
ViT-S: Ablation study shows the effect of the amount of downstream fine-tuning data vs the fine-tuning methods, Test AUC vs % training data for AD classification.

**Figure 3.**
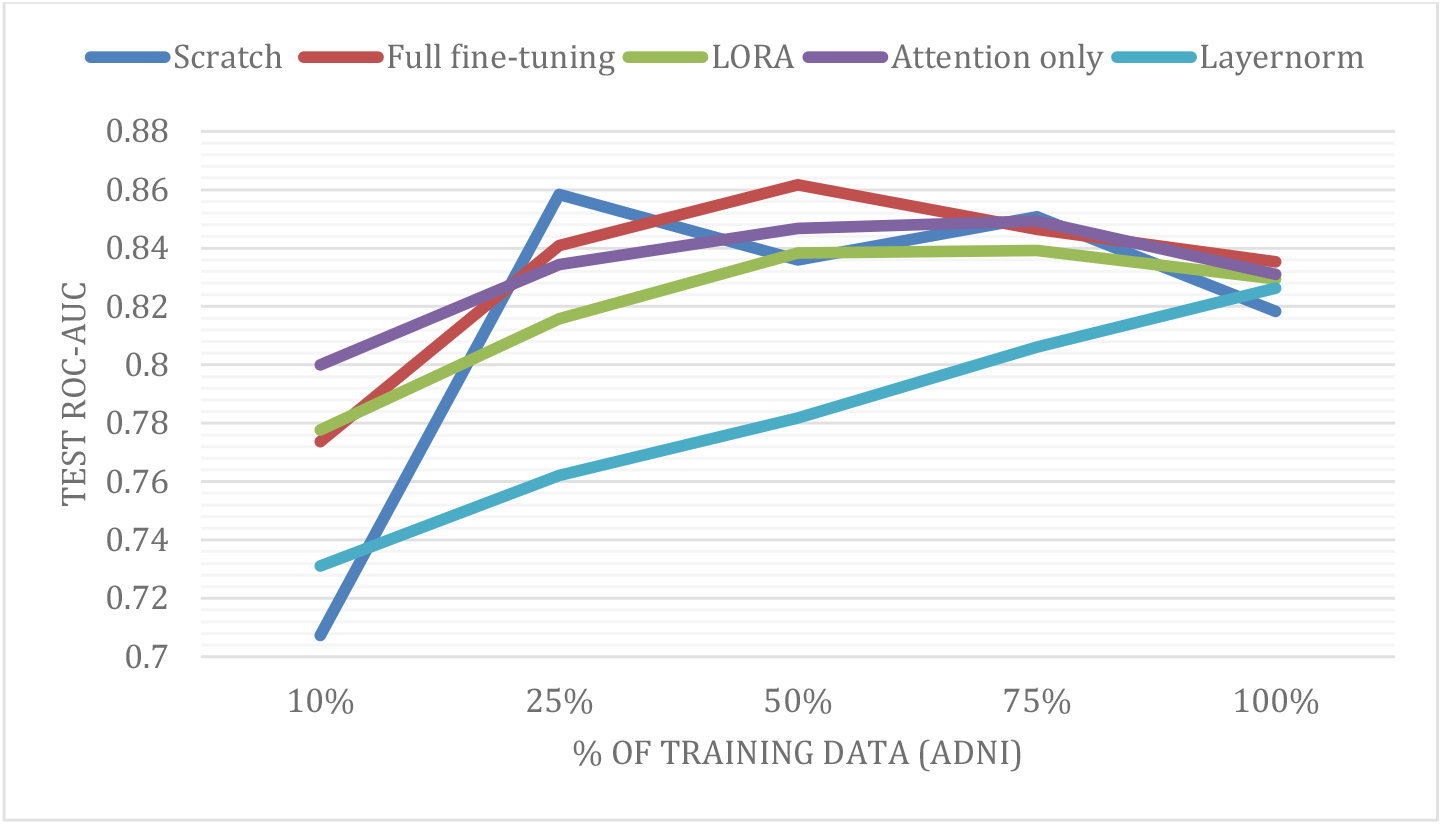
VIT-B: Ablation study shows the effect of the amount of downstream fine-tuning data vs the fine-tuning methods, Test AUC vs % training data for AD classification.

## 5. DISCUSSION

### 1. Test performance for PEFT is competitive with or outperforms full fine-tuning, but at a reduced cost

The PEFT methods reduced the number of trainable parameters significantly compared to full fine-tuning i.e., 30% to 0.04% of the total model size. For AD classification, attention tuning matched the full fine-tune performance on the ADNI test set when trained with 100% of the training data but with a reduced model size – i.e., only using 32% of the original number of model parameters. With LORA (1.3% model parameters) and layernorm fine-tune (0.04% model parameters), these methods give a considerable reduction in model size. We recorded only a slight drop in the test ROC-AUC relative to full fine-tune. Another key takeaway was that all of the PEFT methods were comparable in performance with the full fine-tune on the zero-shot OASIS test set. For brain age prediction, all the PEFT methods outperformed both training from scratch and full fine-tuning. For ViT-B, attention fine-tuning was the best-performing PEFT method achieving a reduction in MAE by 0.42 years. For PD classification, all PEFT methods showed only a slight drop in performance compared to full fine-tune but always outperformed training from scratch. LORA and Layernorm fine-tune outperformed full fine-tune on the zero-shot UPenn dataset. For both the brain age prediction and PD classification tasks, PEFT outperforms the 3D DenseNet121 CNN.

### 2. PEFT methods boosted performance in limited data settings

As shown in **Figure 1**, for ViT-B, the attention fine-tune and LORA methods outperformed (+0.004% to +3% test ROC-AUC) full fine-tuning when using 10% (258 scans) of the ADNI training data set for AD classification. We illustrate the overall effect of the downstream training dataset size (10% to 100%) on the PEFT test performance in **Figure 1**. All PEFT methods tested outperformed training from scratch by 3% to 10% in terms of the test ROC-AUC for AD classification. All the PEFT methods also outperformed the 3D DenseNet 121 CNN for AD classification when using only 258 training scans, including a performance boost of almost 11% with the attention fine-tune method. The PEFT provides the ViT with the capability to be competitive with the 3D DenseNet121 CNN, even with limited training data.

### 3. PEFT helped in adaptation to multiple downstream tasks and datasets

We tested the effectiveness of the PEFT with multiple downstream neuroimaging tasks and multiple datasets as shown in **Table 3**. These tasks were chosen to provide a diverse set of challenges to evaluate the PEFT methods. The PEFT methods were competitive or outperformed full fine-tuning with a substantial reduction in the number of trainable parameters for AD, PD classification and brain age prediction. The PEFT methods also usually outperformed training the ViTs from scratch.

### 4. Smaller models are competitive with larger model performance

We tested the different PEFT methods as a function of the ViT model size and observed that the smaller ViT-S was competitive and sometimes outperformed the larger model in AD classification. This is likely due to the limited size of typical neuroimaging datasets; the approach helps with efficient fine-tuning of foundation models for new tasks. Like the larger ViT (ViT-B), the PEFT methods help boost ViT-S test-time performance and crucially in low-training data regimes as seen in **Figure 2**. Attention-tuning is the best performing PEFT method for ViT-S for AD classification similar to the ViT-B architecture, both relative to full-finetuning.

We plan to extend this work further with other PEFT methods, additional downstream tasks and other model architectures.

## 5. CONCLUSION

In this work, we evaluated multiple parameter-efficient fine-tuning methods for neuroimaging tasks and datasets. Our experiments demonstrated the benefit of parameter-efficient fine-tuning methods applied to pre-trained vision transformer encoders in resource constrained environments. The PEFT methods were competitive with or outperformed full fine-tuning with fewer model parameters. PEFT boosted performance in low training-data settings. We also show that the PEFT methods can be used to adapt pre-trained backbones to a range of tasks. Additionally, we tested multiple model sizes in a series of ablation experiments for each of the PEFT methods.

